# Salt Effects on the Thermodynamics of a Frameshifting RNA Pseudoknot under Tension

**DOI:** 10.1101/048801

**Authors:** Naoto Hori, Natalia A. Denesyuk, D. Thirumalai

**Affiliations:** Biophysics Program, Institute for Physical Science and Technology, University of Maryland, College Park, MD 20742, USA^†^

**Author notes:** Present address: Department of Chemistry, University of Texas, Austin, TX 78712, USA.

## Abstract

Because of the potential link between -1 programmed ribosomal frameshifting and response of a pseudoknot (PK) RNA to force, a number of single molecule pulling experiments have been performed on PKs to decipher the mechanism of programmed ribosomal frameshifting. Motivated in part by these experiments, we performed simulations using a coarse-grained model of RNA to describe the response of a PK over a range of mechanical forces (*f*s) and monovalent salt concentrations (*C*s). The coarse-grained simulations quantitatively reproduce the multistep thermal melting observed in experiments, thus validating our model. The free energy changes obtained in simulations are in excellent agreement with experiments. By varying *f* and *C*, we calculated the phase diagram that shows a sequence of structural transitions, populating distinct intermediate states. As *f* and *C* are changed, the stem-loop tertiary interactions rupture first, followed by unfolding of the 3’-end hairpin (I⇌F). Finally, the 5’-end hairpin unravels, producing an extended state (E⇌I). A theoretical analysis of the phase boundaries shows that the critical force for rupture scales as (log *C*_m_)^*α*^ with *α* = 1 (0.5) for E⇌I (I⇌F) transition. This relation is used to obtain the preferential ion-RNA interaction coefficient, which can be quantitatively measured in single-molecule experiments, as done previously for DNA hairpins. A by-product of our work is the suggestion that the frameshift efficiency is likely determined by the stability of the 5’ end hairpin that the ribosome first encounters during translation.

## INTRODUCTION

The multifarious roles RNA molecules play in controlling a myriad of cellular functions [1] have made it important to understand in quantitative terms their folding [2–6] and how they respond to external stresses [7]. Among them, one of the simplest structural motifs is the RNA pseudoknot (PK), which is involved in many biological functions. The simplest type, referred to as H-type PK, consists of two stems connected by two loops in which one of the stems forms base pairs with the loop of the other. The PK motif, found in many RNA molecules such as telomerase [8], mRNA [9], ribosomal RNA [10], transfer-messenger RNA, and viral RNA [11], is functionally important [12]. For instance, a PK in the human telomerase RNA is essential for enzyme (a ribonucleoprotein complex) activity [13]. Many pathogenic mutations in the human telomerase RNA are found in the well-conserved PK region of the RNA, further underscoring the importance of the PK [8, 13]. The presence of PKs is also important in ribozyme catalysis and inhibition of ribosome translation.

Recently, there is heightened interest in the biophysics of PK folding because it plays a crucial role in affecting the efficiency of −1 programmed ribosomal frameshifting (−1 PRF) [14–16]. Usually, proteins are synthesized when the ribosome reads the mRNA code in three nucleotide steps until the stop codon is reached. In −1 PRF, the open reading frame of mRNA being translated within the ribosome is programmed to be shifted by one nucleotide, and consequently, the mRNA becomes nonfunctional or produces an entirely different protein [5, 17–20]. The downstream PK of the mRNA is a roadblock at the entrance to the mRNA channel on the ribosome, which impedes the translocation of the ribosome. The ribosome must unfold the PK, presumed to occur by exertion of mechanical force, to complete translocation. Because frameshifting efficiency could depend on how the PK responds to force, a number of single-molecule pulling experiments have focused on PKs [15, 21, 22]. Several factors could determine −1 PRF, as evidenced by the multitude of proposals based on many experimental studies [14, 15, 23–28]. Nevertheless, given that ribosome translocation exerts mechanical force on the downstream mRNA, it is physically reasonable to assume that resistance of PK to force should at least be one factor determining the frameshifting efficiency [14, 29].

The considerations given above and the availability of both ensemble and single-molecule pulling experimental data prompted us to focus on the effects of temperature, mechanical force, and salt concentration on a 28nucleotide H-type RNA PK from the beet western yellow virus (BWYV) [19]. This PK has been the subject of several ensemble experiments, which have characterized the folding thermodynamics as a function of monovalent and divalent cations [30, 31]. The BWYV PK has two stems (S1 and S2) connected by two loops (L1 and L2) as shown in Fig. 1.. The crystal structure reveals that L2 forms triplex-type tertiary interactions with S1[32]. In another context, it has been shown that such interactions and mutations affecting them dramatically change the efficiency of frameshifting [21]. In *vitro* and *in vivo* experiments on many mutants of BWYV suggest variations in the frameshift efficiency, which, for some mutants, is attributed to changes in the stem-loop interactions affecting the stability of PK. For others, direct involvement of the ribosome is invoked [25]. In addition to mechanical force (*f*), it is known from several ensemble experiments that the stability of BWYV PK, like other RNA molecules, depends on monovalent and divalent cations [33]. Thus, it is important to understand in quantitative detail how this crucial structural motif respond to *f* and changes in ion concentration.

**FIG. 1.**
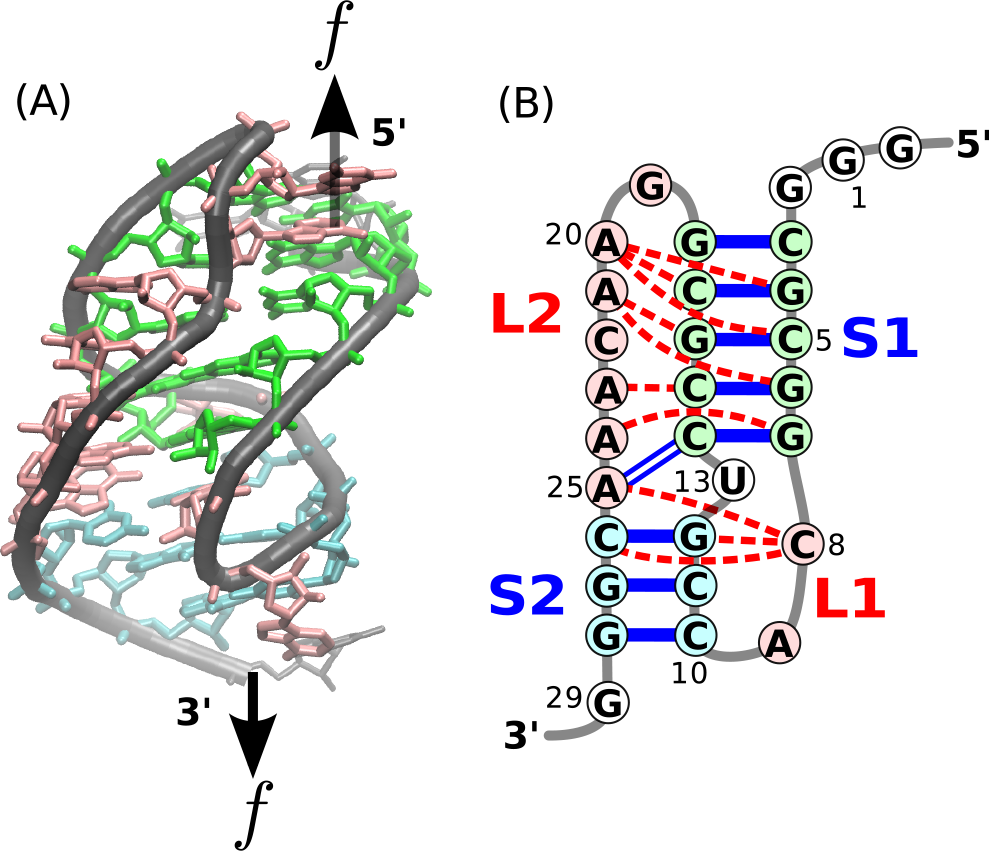
Structure of the BWYV pseudoknot. (**A**) Sketch of the crystal structure [32] color-coded according to secondary structures. The black arrows indicate positions at which the external mechanical force is applied. (**B**) Schematic representation of the secondary structure. Unlike the typical H-type pseudoknot, the two helix stems (S1 and S2) are not coaxially stacked in the BWYV pseudoknot [25]. Tertiary interactions are indicated by red dashed lines.

Although single-molecule pulling experiments suggest how force affects the stability of RNA [19], structural information obtained from such experiments is limited only to changes in molecular extensions. In order to provide a theoretical basis for the changes in the stability and structural transitions as *f* and monovalent salt concentration (*C*) are altered, we performed simulations using a coarse-grained three-site-interaction model (TIS model) [34] of the PK. After demonstrating that our simulations reproduce the temperature-induced structural transitions observed in ensemble experiments, we calculated the global phase diagrams in the [*C*, *f*] plane. The simulations, performed in the same force range as used in Laser Optical Tweezer (LOT) experiments [19], quantitatively reproduce the experimental observations. By combining our simulations and a theory based on ion-RNA interactions expressed in terms of preferential interaction coefficients [31, 35], we produce several key predictions: (1) The phase diagram in the [*C*, *f*] plane shows that the folded state (F) ruptures in stages populating distinct intermediates as *f* increases. Structurally, the low force intermediate is similar to the one populated during thermal melting. The sequence of *f*-induced structural transitions can be described as F⇌I⇌E, where E is an extended state; (2) The critical force, *f*_c_, to rupture the PK scales as (log *C*_m_)^*α*^ with *α* = 1 for the I⇌E transition and *α* = 0.5 for the I⇌F transition. We expect this result to be valid for H-type PKs composed of stems with vastly different stability, such as the BWYV PK. This result is of general validity and is applicable to other PKs as well. The slope of the *f*_c_ versus log *C*_m_ is proportional to the changes in the preferential interaction coefficient between an intermediate and unfolded states. This prediction can be used in conjunction with single-molecule experiments allowing for direct measurement of a key quantity in ion-RNA interactions; and (3) To the extent the downstream PK is a roadblock for translation, our work suggests that the ribosome should exert a force ~ (10 – 15) pN [36] for unimpeded translocation along mRNA, although the actual force value could vary depending on factors such as interaction between PK and ribosome. Based in part on our work we propose that there should be a link between the stimulatory effect of frameshifting and the stability of the 5’–end of the PK. Our predictions are amenable to validations using single-molecule experiments.

## RESULTS AND DISCUSSION

### Temperature-induced transitions in BWYV PK

In order to validate our model, we first performed temperature replica-exchange molecular dynamics (REMD) simulations at *C* = 500 mM with *f* = 0. Formations of stems (S1 and S2) and tertiary (stem-loop) interactions (L1 and L2) are characterized by hydrogen bond (HB) energies 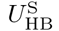 and 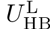, respectively. Changes in the distributions of HB energies as the temperature is increased are shown in Fig. S1 in the Supporting Information. (Fig. S1 shows that there are two minima corresponding to two distinct states. The minimum that dominates at *T* = 20°C corresponds to the F state in which both S1 and S2 are formed (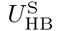 ≈ −45 kcal/mol). The minimum with 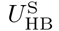 ≈ −30 kcal/mol at 80°C corresponds to a state in which S1 is intact (Fig. S1). The free energy profile of 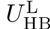 shows only one possible minimum around 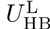 ≈ −20 kcal/mol at *T* = 20°C, indicating that L1 and L2 fold/unfold cooperatively as temperature changes.

We calculated the free energy profiles *G*(*R_*α*_* = −*k*_B_*T*log*P*(*R*_*α*_ (R_*α*_ = *R*_ee_, end-to-end distance, or *R*_g_, radius of gyration) from the distributions of *P*(*R*_*α*_), which are given in Fig. S2 in the Supporting Information. The locations and the number of minima report on the thermodynamic transitions between the states (Fig. 2). (1) At a low temperature, T = 20°C, the PK is completely folded (F state) including all the structural motifs, S1, S2, L1 and L2. The existence of F state is also confirmed by the presence of a minimum at *R*_ee_ ≈ 3 nm (Fig. 2A) and at *R*_g_ ≈ 1.3 nm (Figure 2B). These values are similar to those obtained directly from the crystal structure (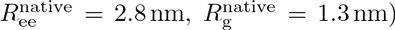). (2) At *T* = 60°C, the free energy profile for 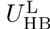 shows that some of the tertiary interactions, involving L1 and L2, are no longer stably formed (Fig. S1B). On the other hand, *G*(*R*_g_) shows two stable minima, indicating that there are two possible states ((Fig. 2B). In one state, both stems S1 and S2 are formed but tertiary interactions are absent, corresponding to the I1 state identified using UV absorbance and differential scanning calorimetry experiments (Fig. 3) [30, 31]. The other conformation, in which only S1 is formed, has been also observed in experiments and is referred to as the I2 state. It should be noted that the distribution of 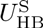 also has two minima at 60°C (Fig. S1A) whereas *G*(*R*_ee_) has only one minimum (Fig. 2A). (3) At a still higher temperature *T* = 80°C, *G*(*R*_ee_) and *G*(*R*_g_) each have a minimum with S1 stably formed. This is also reflected in (Fig. S1, which shows a minimum at 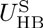 ≈ −30 kcal/mol and a minimum at 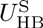 ≈ 0. Thus, completely unfolded conformations, U state, and the I2 state are populated. (4) At *T* = 110°C, both 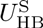 and 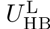 are 0, indicating that only the U state exists. This is also reflected in *G*(*R*_ee_) and *G*(*R*_g_), which show that the PK is completely unfolded. In both profiles (Fig. 2A and Fig. 2B), the center of minimum is located at larger values than in the F state (*R*_ee_ ≈ 5 nm and *R*_g_ ≈ 2 nm).

**Fig. 2.**
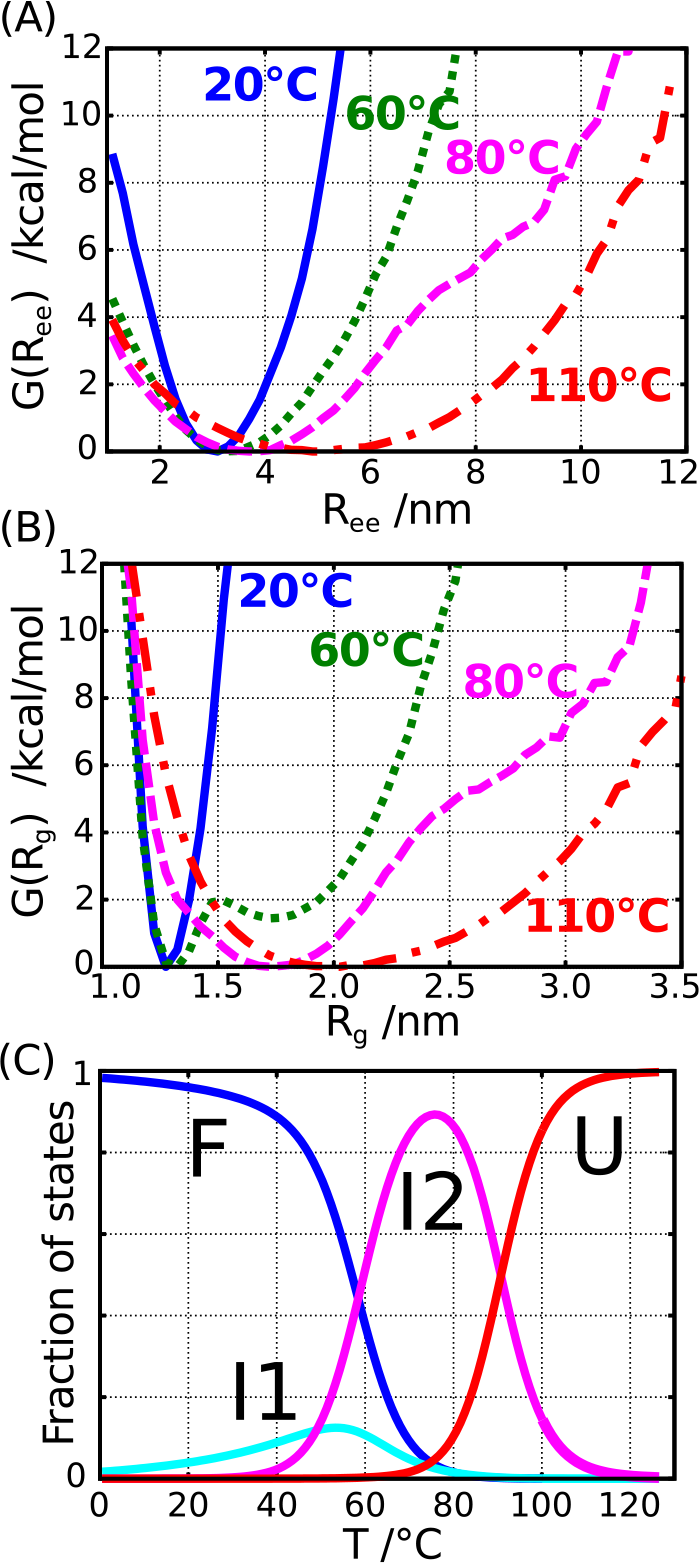
Characteristics of the thermally induced folding-unfolding transition of BWYV PK simulated using the temperature-REMD method at C = 500 mM and *f* = 0.(A and B) Free energy profles of the molecular extension (A;*R*_ee_) and radius of gyration (B; *R*_g_) at 20, 60, 80, and 110°C.Each curve is normalized to G = 0 at its probability *P*(*R*) maximum. (C) Temperature dependence of the fraction of the four prominent states, which are explicitly labeled.

The simulation results in Fig. 2 and Fig. S1 show that melting of BWYV PK undergoes through two intermediate states, F⇌I1⇌I2⇌U where I1 has the topology of the F state but without significant tertiary interactions (Fig. 3). Although simulations can be used to distinguish I1 and F states, the intermediate does not have significant population. The identification of I1 in our simulations provides a structural support to experiments in which I1 was inferred from three broad overlapping peaks in the heat capacity curves [30, 31]. It should be stressed that the experimental heat capacity curves exhibit only two discernible peaks, which would normally imply that there is only one intermediate present. The range of melting temperatures for the three transitions can be obtained from the temperature dependence of the various states *f*_*α*_(*T*) (*α*=F, I1, I2 or U) given in fig. 2C. The I1 state has a maximum population around *T* ≈ 50°C. We equate this with the melting temperature *T*_m1_ associated with the F⇌I1 transition. Similarly, *T*_m2_ for the I1⇌I2 transition is ≈ 70°C. The *T*_m3_ for the I2⇌U transition occurs around 90°C (*f*_I2_(*T*_m3_) = *f*_u_(*T*_m3_)). The values of the three melting temperatures obtained in our simulations are in remarkable agreement with experiments 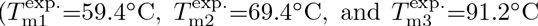 taken from results of a differential scanning calorimetry experiment at pH=7 in Table 1 of Nixon and Giedroc [30]).

**Fig. 3.**
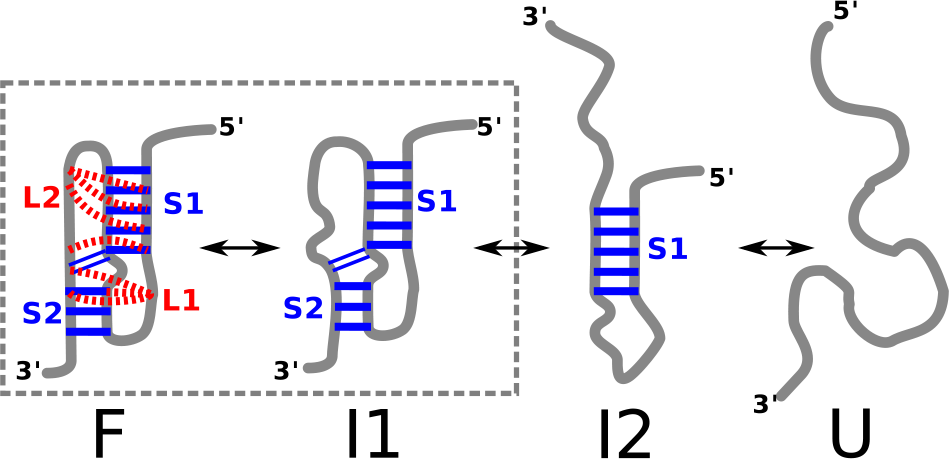
Schematic of the thermodynamic assembly of BWYV pseudoknot inferred from simulations at zero force. Besides folded (F) and unfolded (U) states, there are two intermediate state, I1 and I2. The topology of the I1 state is the same as F, except tertiary interactions are absent.

The dependence of the stability of the F state with respect to the U state, Δ*G*_UF_(*T*) is calculated using a procedure that does not rely on any order parameter [37].The value of Δ*G*_UF_(T = 37°C) is −14.3 kcal/mol from simulations, which is in excellent agreement with experiments. The experimental value reported by Nixon and Giedroc for Δ*G*_UF_(T = 37°C) = −13.3 kcal/mol, whereas Soto et al. estimated that Δ*G*_UF_(*T* = 37°C) = −15.1 kcal/mol [30, 31]. We also predict that Δ*G*_UF_(*T* = 25°C) is −19.0 kcal/mol. The excellent agreement between simulations and experiments for the temperature induced structural transitions and the Δ*G*_UF_(*T*) validates the model allowing us to predict the effect of *f* on BWYV PK.

**Diagram of states in the** [*C*, *f*] **plane**

In order to obtain the phase diagram in the [*C*, *f*] plane of the BWYV PK, we performed a series of low friction Langevin dynamics simulations by varying the salt concentration from 5 to 1200 mM and the mechanical force from 0 to 20 pN, at a constant temperature of 50°C. We varied *C* by changing the Debye length in the simulations, and the mechanical force was externally applied to the terminal nucleotides (Fig. 1A).

Determination of the phase diagram in the [*C*, *f*] plane requires an appropriate order parameter that can distinguish between the various states. In single-molecule pulling experiments, the variable that can be measured is the molecular extension, *R*_ee_, which is conjugated to the applied force. The overall size of the RNA molecule is assessed using the radius of gyration, *R*_g_. Therefore, we characterized the states of the RNA using both *R*_g_ and *R*_ee_ as order parameters. The values of the predicted *R*_g_ at f = 0 can be measured using scattering experiments. Using these two parameters, we obtained the [*C*, *f*] phase diagram ((Fig. 4). Comparison of the diagram of states in (Fig. 4D and (Fig. 4E reveals common features and some differences. From (Fig. 4D and (Fig. 4E, we infer that at f > 12.5 pN, extended (E) conformations are dominant at all values of *C*. As the force decreases, the PK forms compact structures. The boundary separating the extended and relatively compact phases depends on the salt concentration and the value of the mechanical force. The critical force to rupture the compact structures increases linearly as a function of logarithm of salt concentration (Fig. 4D; boundary between red and green regions). At low forces (*f* < 2.5 pN), the diagram of states based on *R*_ee_ shows that the extension is relatively small as *C* changes from a low to a high value. From this finding, one might be tempted to infer that the PK is always folded, which is not physical especially at low (*C* ≈ 10 mM) ion concentrations. In contrast, (Fig. 4E shows that below 5 pN, there is a transition from compact structures (*R*_g_ ≈ 1.3 nm in the blue region) at high *C* to an intermediate state (*R*_g_ > 2.2 nm in the green region) at C ≈ 100 mM. The differences between the diagrams of states in the [*C*, *f*] plane using *R*_ee_ and *Rg* as order parameters are more vividly illustrated in terms of the free energy profiles *G*(*R*_*α*_) = −*k*_B_*T*log(*R*_*α*_) where *R*_*α*_ is *R*_ee_ or *R*_g_ (Fig. 5). The profiles *G*(*R*_ee_), at three values of *C* and different *f* values, show two minima at most. At *f* = 0, there is only one discernible minimum at *R*_ee_ ≈ 4.2 nm at C = 10 mM. The minimum moves to *R*_ee_ ≈ 3 nm at *C* = 1200 mM corresponding to a folded PK. At *f* = 5 pN there are two minima at *C* = 10 mM corresponding to a compact structure and a stretched conformation (see the cyan profile in (Fig. 5A). As *f* exceeds 5 pN, there is essentially one minimum whose position increases as *f* increases. In the force regime (*f* > 5 pN), only the E state is visible at all *C*.

**Fig. 4.**
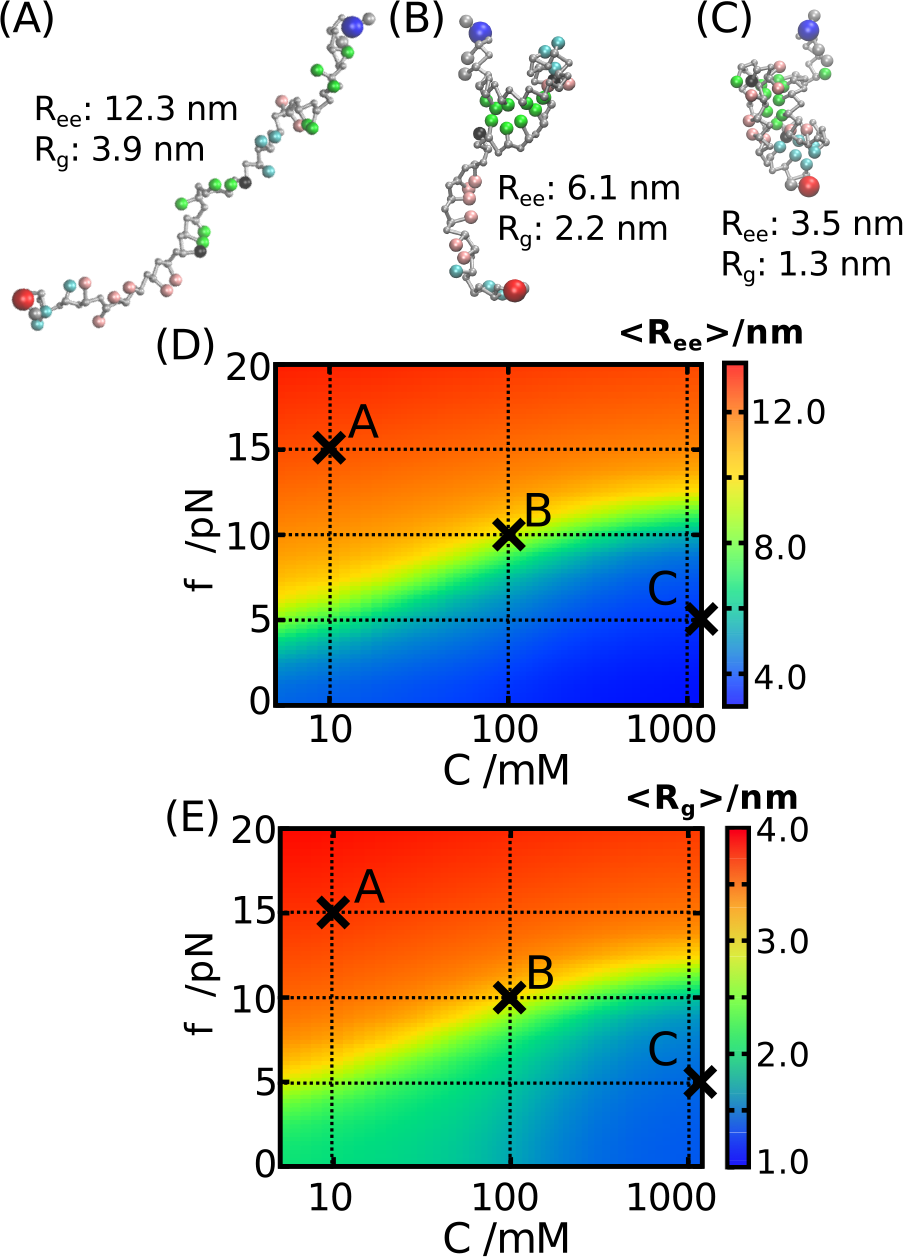
(**A**–**C**) Representative snapshots corresponding to the three distinct states from simulations. (**D**) Diagram of states in the [*C*, *f*] plane obtained using extension *R*_ee_ as the order parameter. (**E**) Same as D, except *R*_g_ is used to distinguish the states. The three crosses in D and E correspond to conditions from which the snapshots (*A*-*C*) are sampled. The scale for *R*_ee_ and *R*_g_ are given on the right.

**Fig. 5.**
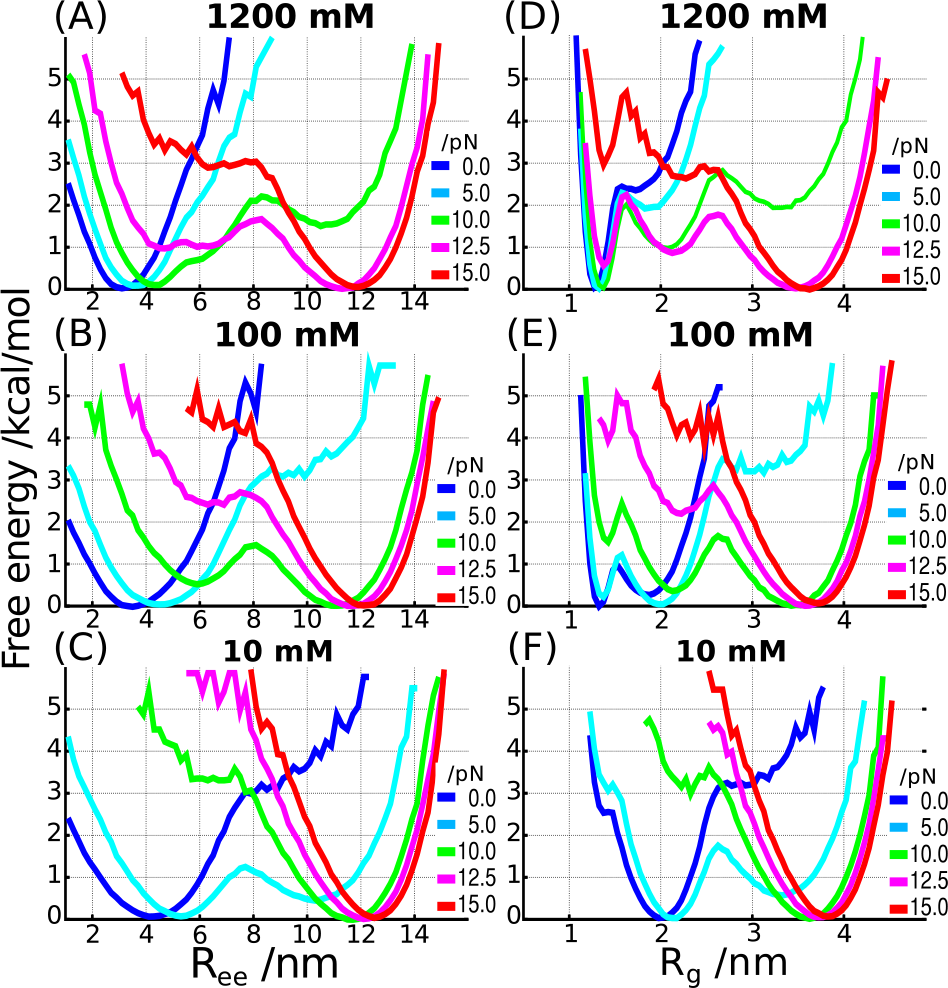
Free energy profiles as a function of *R*_ee_ (**A**-**C**) and *R*_g_ (**D**-**E**) at three different salt concentrations and a range of forces. The locations of the minima change as *C* and *f* vary, indicating transitions between distinct states of the BWYV PK.

In order to compare the thermal free energy profiles generated at *C* = 500 mM (Fig. 2), we calculated the *G*(*R*_g_) and *G*(*R*_ee_) at different forces with *C* fixed at 500 mM. Comparison of (Fig. 2 and (Fig. S3 shows that at *f* = 0 the free energy profiles are similar. At *f* ≠ 0 the folded state is destabilized, and at *f* = 15 pN, the unfolded state is more stable than the folded PK.

A much richer diagram of states emerges when the *G*(*R*_g_) = −*k*_B_*T*log*P*(*R*_g_) is examined. The *G*(*R*_g_) profile at *C* = 10 mM is similar to *G*(*R*_ee_). However, at higher concentrations, there is clear signature of three states (especially at *f* = 12.5 pN) corresponding to the folded PK, an intermediate state, and the E state. The free energy profiles show that under force, there are minimally three states, which are F, I, and E. Because *R*_ee_ cannot capture the complexity of the states populated as *C* and *f* are varied, it is important to consider *R*_g_ as well to fully capture the phase diagram of the PK. We also computed the free energy profiles as a function of root-mean-square-deviations (RMSD). In terms of the number and relative positions of basins, the profiles based on RMSD are qualitatively similar to one of *R*_g_ (Fig. S4).

### Formation of stems and loops

In order to highlight the structural changes that occur as *C* and *f* are varied, we used, *Q*, the fraction of native HB interactions as an order parameter. The [*C*, *f*] phase diagram (Fig. 6A) calculated using the average *Q* is similar to the *R*_g_-phase diagram (Fig. 4E), indicating the presence of three states (Fig. 6A). Using *Q*, we can also quantitatively assess the contributions from different structural elements to the stability of the PK and correctly infer the order in which the structural elements of the PK rupture as *f* is increased. In order to determine the structural details of each state, we calculated the individual *Q*s for the two stems (*Q*_s1_, *Q*_S2_) and the two loops (*Q*_L1_ *Q*_l2_) (Fig. 6B). The dependence of (*Q*_S1_) as a function of *C* and *f* shows that Stem 1 (S1) is extremely stable and remains intact at all salt concentrations, rupturing only at f ≈ 10 pN (Fig. 6B, upper left panel). In contrast, the upper right panel in Fig. 6B shows that Stem 2 (S2) is only stably folded above a moderate salt concentration (*C* ≈ 80 mM) and low *f*. The stability difference between S1 and S2 can be explained by the number of canonical G-C base pairs; S1 has five G-C base pairs, whereas only three such pairs are in S2. Consequently, S1 cannot be destabilized even at low *C*, but its rupture to produce extended states requires the use of mechanical force. Above *C* > 100 mM, the fraction of contacts associated with S2 and the two loops (*Q*_S2_, *Q*_L1_ and *Q*_L2_) simultaneously increase. All the interactions are, however, still not completely formed (*Q* ≈ 0.8) even at the highest salt concentration, *C* = 1200 mM. In particular, the contacts involving L2, *Q*_L2_ ≈ 0.6, implying that tertiary interactions are only marginally stable at *T* = 50° C.

**Fig. 6.**
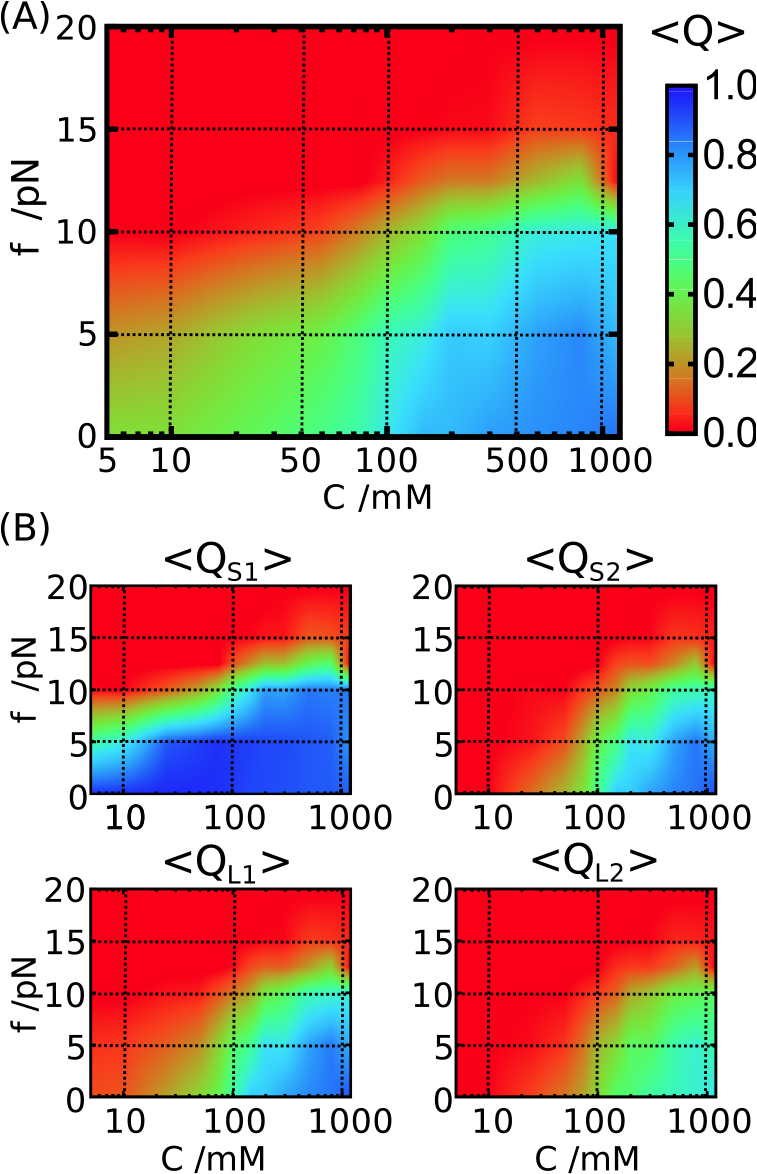
[*C*, *f*] phase diagram using the fraction of native contacts as the order parameter. **(A)** The diagram of states determined using an average of the total *Q* for the PK. The scale is given on the right. **(B)** Decomposition of *Q* into stems and loops: S1, stem 1; S2, stem 2; L1, loop 1; L2, loop 2.

The results in Fig. 6 provide a structural explanation of the transitions that take place as *C* and *f* are changed. (1) In the high force regime, the two stems are fully ruptured and native interactions are absent creating the E state. (2) Below *C* ≈ 100 mM and *f* ≲ 10 pN (left bottom on the [*C*, *f*] plane), only S1 is intact, and thus, a hairpin conformation is dominant. This is the intermediate state, I2, identified by Soto et al. who found that S1 remains stable even at salt concentration as less as 20 mM NaCl [31]. (3) Above *C* ≳ 100 mM and *f* ≲ 10 pN (right bottom), both S1 and S2 stems are formed, and the tertiary interactions involving the two loops are at least partly formed, ensuring that the topology of the PK is native-like. At *f* = 0 and temperature as a perturbation, it has been shown that another intermediate state, I1, exists between I2 and folded states [30, 31]. The I1 state may correspond to conformations in which two stems are fully folded with the entire topology of the native conformation intact, but tertiary interactions between stems and loops are incomplete. Although we have accounted for the presence of such a state in our simulations (see Fig. 2), from here on, we do not distinguish between I1 and the completely folded state in the phase diagram since the increases in *Q*_S2_, *Q*_l1_ and *Q*_L2_ on the [*C*, *f*] plane almost overlap. Therefore, we classify the right bottom region in the [*C*, *f*] phase diagram as (I1 + F) state.

### Population analysis of the secondary and tertiary interactions

The [*C*, *f*] phase diagrams provide a thermodynamic basis for predicting the structural changes that occur as *f* and *C* are varied. However, they capture only the average property of the ensemble within each state without the benefit of providing a molecular picture of the distinct states, which can only be described using simulations done at force values close to those used in LOT experiments. Therefore, we investigated the distribution of interactions to ascertain the factors that stabilize the three states as *C* and *f* are varied. Our analysis is based on the HB energy, *U*_HB_. The energy function for HB captures the formation of base pairs and loop-stem tertiary interactions, taking into account both distance and angular positions of the donor and the acceptor [37].

Fig. 7 shows that the probability densities of the total HB energy have three peaks at *U*_HB_ ≈ 0, −30, and −60 kcal/mol. In the high-force regime (*f* > 12.5 pN), there is only one population at all *C* around *U*_HB_ ≈ 0, which obviously corresponds to the fully extended state. The total HB energy can be decomposed into contributions arising from four distinct structural motifs in the PK (Fig. S5). The peak at *U*_HB_ ≈ −30 kcal/mol (Fig. 7) arises due to the formation of S1 (compare Figs. 7 and S5A). The broad peak at *U*_HB_ ≈ −60 kcal/mol (Fig. 7A) is due to the sum of contributions from S2 and the tertiary interactions.

**Fig. 7.**
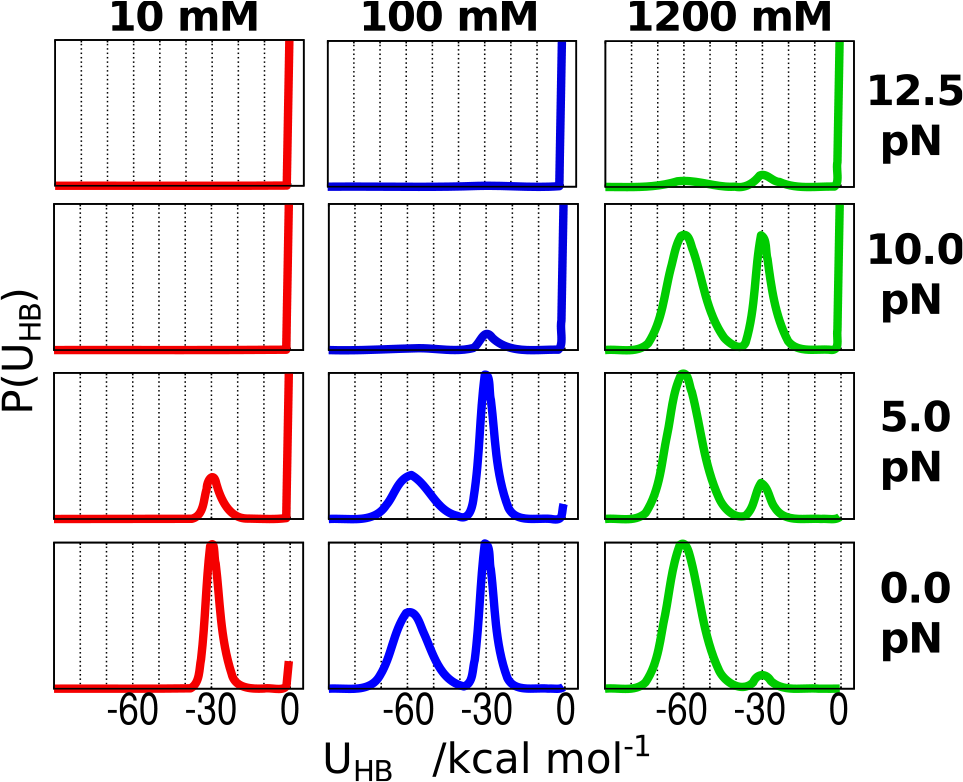
Probability distributions of the total hydrogen bond energy as a function of *C* and *f*, which are labeled. The distinct peaks are fingerprints of the states that are populated at a specified *C* and *f*.

Parsing the *U*_HB_ due to contributions from interactions among S1, S2, L1, and L2 produces a picture of thermodynamic ordering as *C* (*f*) is increased (decreased) from low (high) value to high (low) value. Low *C* and high *f* correspond to the top left region in the phase diagrams (Figs. 4 and 6). In all conditions, S1 is the first to form as indicated by the presence of distinct peaks at *C* > 100 mM and f ≲ 10 pN in Fig. S5A. Under more favorable conditions (*f* < 10 pN; *C* ≈ 300 mM or greater) S2 and tertiary interactions involving L1 are established (Figs. S5 B and C). Only at the highest value of *C* and *f* ≲ 5 pN, tertiary interactions involving L2 emerge. Interestingly, the order of formation of the various structural elements en route to the formation of their folded state follows the stabilities of the individual elements with the most stable motif forming first. The PK tertiary interactions are consolidated only after the assembly of the stable structural elements. This finding follows from our earlier work establishing that the hierarchical assembly of RNA PKs is determined by the stabilities of individual structural elements [38].

We did not find any condition in which S1 and S2 are completely folded, but there are no tertiary interactions, 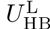 ≈ 0 (i.e., isolated I1 state). Our simulations show that the tertiary interactions contribute to the loop energy only after the stems are formed. Thus, the actual population is distributed between I1 and F, and the tertiary interactions are only marginally stable in the conditions examined here. Soto et al. suggested that diffuse Mg^2+^ ions play an important role in stabilizing such tertiary interactions [31]. Because Mg^2+^ is not considered here, we cannot determine the precise interactions stabilizing the I1 state, especially at *f* ≠ 0. In the discussion hereafter, for simplicity, we refer to (I1 + F) state as F.

### Critical rupture force and preferential ion interaction coefficient

In order to delineate the phase boundaries quantitatively, we calculated the free energy differences between the three major states in the [*C*, *f*] plane using 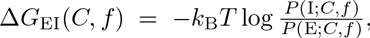, where the classification of the states, E, I (=I2), or F, is based on the HB interaction energies of stems, 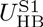 and 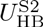 (Fig. 8). A similar equation is used to calculate Δ*G*_IF_. Based on the distribution of *U*_HB_, we determined the threshold values for S1 formation as 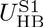 < −15kcal/mol (Fig. S5A). Similarly, S2 formation is deemed to occur if 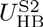 −10kcal/mol (Fig. S5B). We classified each structure depending on whether both stems are formed (F) with both 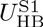 < −15 kcal/mol and 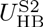 < −10 kcal/mol, only S1 is formed (I (=I2)), or fully extended (E).

**Fig. 8.**
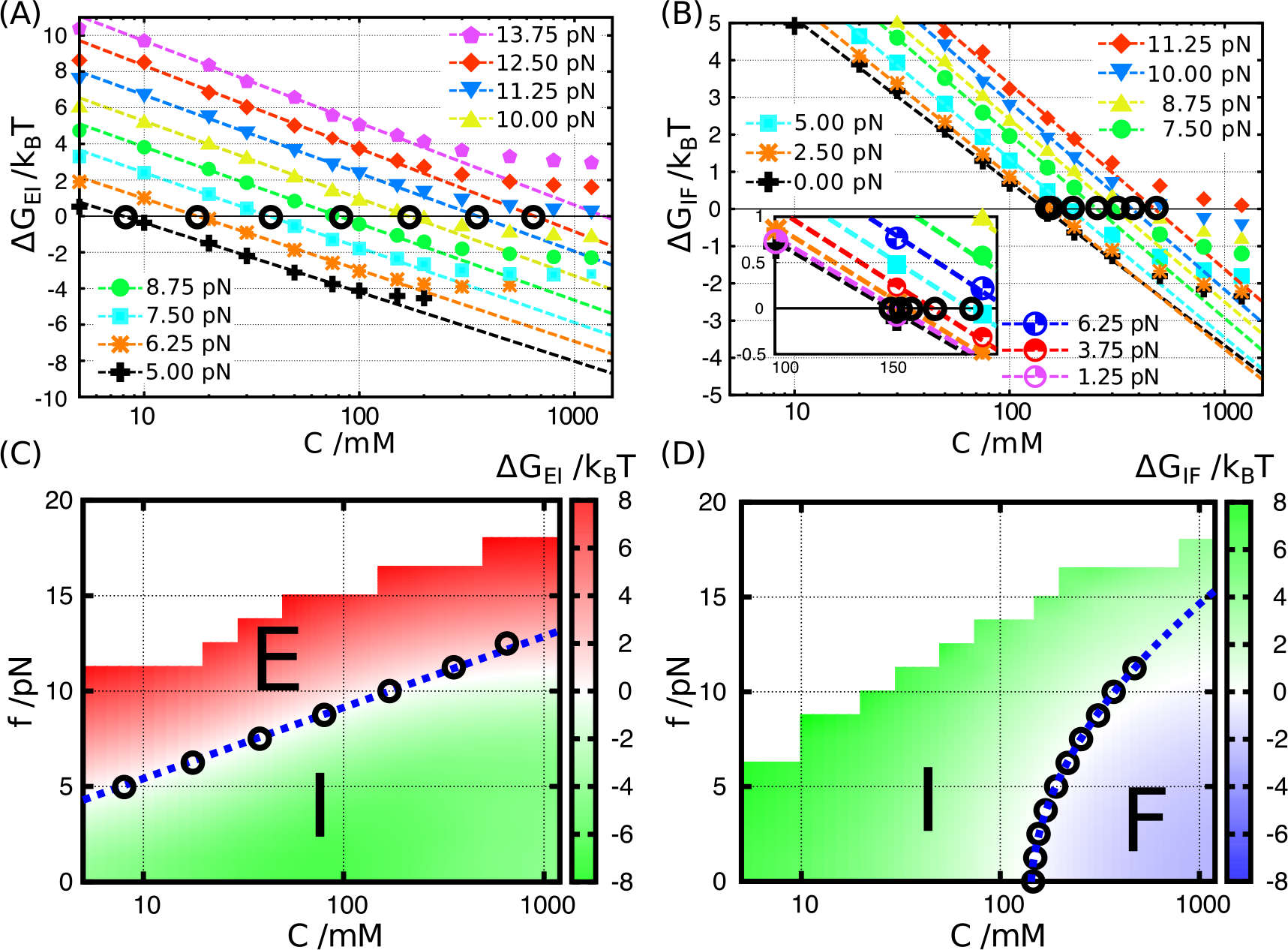
Free energy differences between E and I, and I and F calculated from REMD simulations. The definition of states, E,I, and F, is based on the hydrogen bond interaction energies associated with each structural element (see main text for details).The I state is analogous to I2 in Fig. 3. **(A and B)** Salt concentration dependences of Δ*G* is plotted for different force values (symbols are explained in A and B, respectively). **(C)** Δ*G*_EI_ is shown on the [*C*; *f*] plane, and phase boundary is quantitatively described (white zone between red and green regions). The blue dashed line is the theoretical prediction where *f*_c_ ∝ log *C*_m_ [Equation 4]. Circles (o) indicate conditions with Δ*G*_EI_ = 0 extracted from linear dependence of Δ*G* on log *C* in (A). It should be noted that the E state is under tension, and hence is different from the unfolded state upon thermal melting at *f* = 0: (D) Same as (C) except the results describe I⇌F transition. Here, the critical force describing the phase boundary is a nonlinear function of log *C*_m_ [Eq. 6].

The classification of the diagram of states based on the calculation of Δ*G*_EI_ and Δ*G*_IF_ is consistent with the *R*_g_ and *Q* phase diagrams (Figs. 8C and 8D). At all values of *f*, Δ*G* depends linearly on log*C* over a broad range of *C* as shown in Figs. 8A and 8B. The midpoints of salt concentration, *C*_m_s, at each *f*, determined using 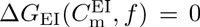 and 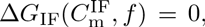, leads to 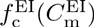 and 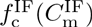, respectively, for the E⇌I and I⇌F transitions. The forces 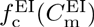 and 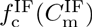 are the critical forces needed to rupture the I and F states, respectively. The locus of points 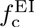 and 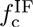(*C*_m_ are shown as circles in Figs. 8C and 8D, respectively. It is clear that 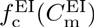 is linear in log 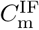 along the phase boundary. In contrast, the dependence of 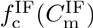 on log 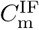 is nonlinear. In what follows, we provide a theoretical interpretation of these results, which automatically shows that the difference in preferential ion interaction coefficients between states can be directly measured in LOT experiments [39–43].

The salt concentration dependence of RNA stability can be determined using the preferential interaction coefficient, defined as 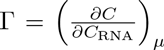, where *C*_RNA_ is the concentration of RNA, and the chemical potential of the ions, μ, is a constant [35, 44]. The free energy difference, Δ*G*_*α*β_(=*G*_β_−*G*_*α*_), between two states α and β (such as E and I or I and F) can be expressed as:
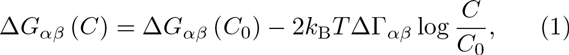
where *C*_0_ is an arbitrary reference value of the salt concentration [45]. Note that we consider only 1:1 monovalent salt such as KCl or NaCl for which ion activity can be related to salt concentration *C*. The factor of 2 in (Eq. 1 arises from charge neutrality. The difference, ΔΓ_*α*β_(=Γ−Γ_*α*_, is interpreted as an effective number of bound or excluded ions when the RNA molecule changes its conformation between the two states [35, 44]. The free energy change upon application of an external force *f* is 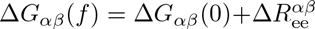 where 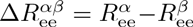[29]. Thus,

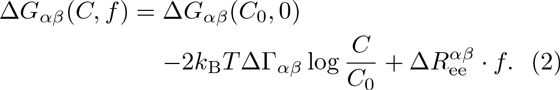

In the [*C*, *f*] phase diagram, Δ*G*_*α*β_ is 0 along the phase boundaries. Since the reference salt concentration *C*_0_ is arbitrary, we determined its value using Δ*G*_*α*β_(*C*_o_, *f* = 0) = 0. Thus, 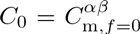 is the salt concentration at the midpoint of the transition at zero force. The determination of ΔΓ_*α*β_ using this procedure from single-molecule pulling data has been described in several key papers [39–42]. By adopting the procedure outlined in these studies and with our choice of *C*_0_, we rearrange (Eq. 2 to obtain the phase boundary at f ≠ 0,
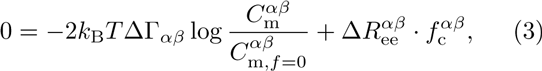
where 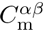 is the midpoint of salt concentration, and 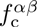 is the critical force associated with the transition between states *α* and β. Measurement of 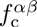 using data from single-molecule pulling experiments has been used to obtain ΔΓ_*α*β_ for few systems that exhibit two-state transitions [39‐42]. Here, we have adopted it for a PK exhibiting more complex unfolding transitions. The connection between the relation in Eq. 3, derived elsewhere, to Clausius-Clapeyron equation was established recently by Saleh and coworkers [41].

It follows from Eq. 3 that the critical force, leading to the prediction that there should be a linear dependence between 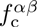 and the logarithm of the midpoint of the salt concentration if 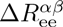 is a constant value along the phase boundary. In accord with the prediction in (Eq. 4), we find that 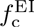 varies linearly with 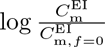 along the phase boundary separating E and I (Fig. 8C) where (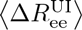) ≈ 4.9nm over a broad range of *f*_c_ (Figs. 9A and 9C).

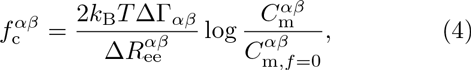

**Fig. 9.**
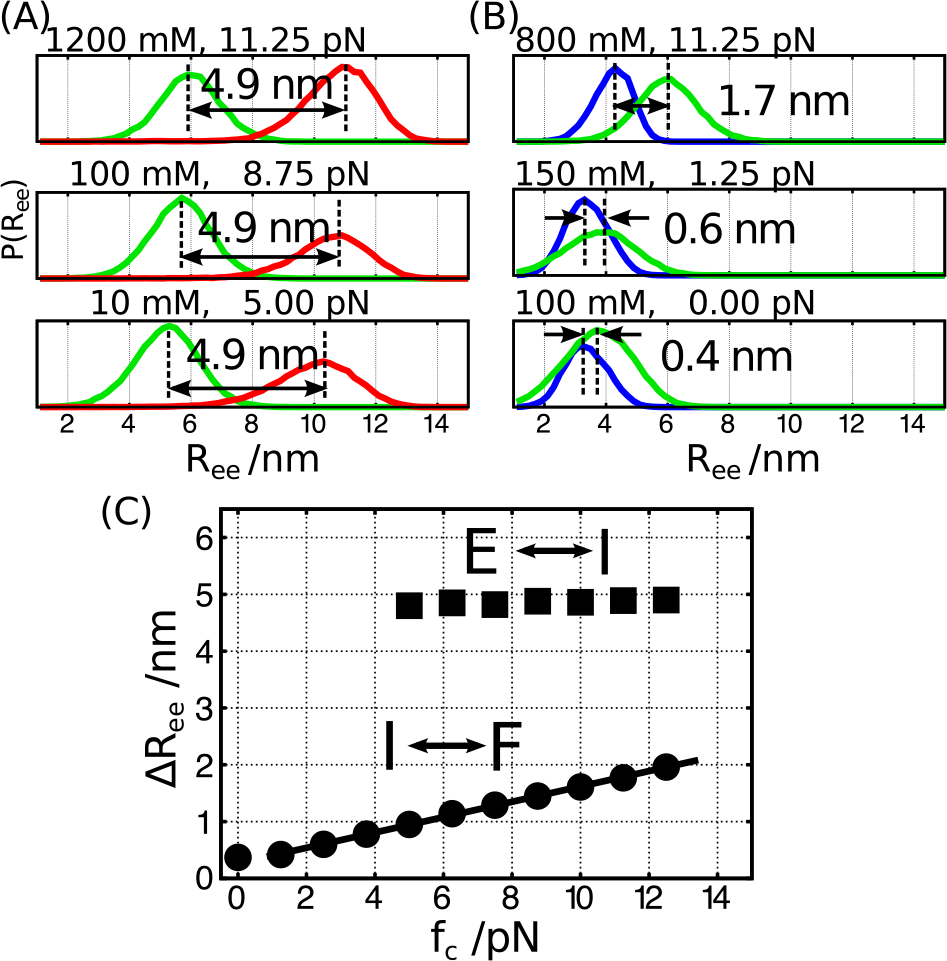
**(A and B)** Probability distributions of the extensions for states E and I (A), and I and F (B). All the conditions shown here are near the phase boundaries in the [*C*, *f*] plane (Δ*G*_EI_ and Δ*G*_IF_ between the two states are close to 0). Distances between peaks of the two distributions are shown. In the E⇌I transition (A), the mean distances are almost constant around 4.9 nm. On the other hand, in the I⇌F transition (B), the mean distance becomes smaller as the mechanical force weakens. **(C)** Dependence of Δ*R*_ee_ on *f*_c_ along the phase boundaries. In the I⇌F transition, the dependence can be fit using 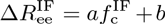, where a = 0.14 nm/pN and b = 0.27 nm (solid line).

The observed nonlinearity of the phase boundary separating I and F over the entire concentration range (Fig. 8D) arises because 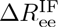 is not a constant but varies linearly with 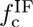 (Figs. 9B and 9C). In giving from F↔I, there is only a rupture of few tertiary interactions and unfolding of the relatively unstable S2. Because this transition is not cooperative (interactions break in a sequential manner), we believe that a linear behavior over a limited force range is not unreasonable. In contrast, the unfolding of S1 is cooperative, like some hairpins studied using pulling experiments, in the E↔I transition, resulting in the constant value for Δ*R*_ee_ over the force range probed. Thus, using 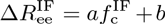 and (Eq. 4, we find that 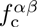 satisfies, 
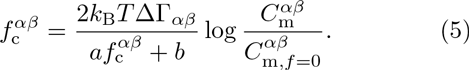
Note that if *a* is zero, then Eq. 5 reduces to Eq. 4 with *b* = 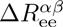 = *const*. Solving Eq. 5, the nonlinear dependence of 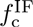 is expressed as,
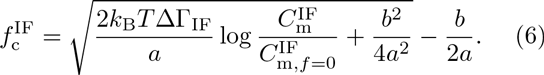
The simulation data can be quantitatively fit using (Eq. 6) (Fig. 8D). In general, we expect 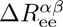depends on 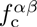, and hence, 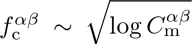. It is worth noting that the estimation of ΔΓ_*α*β_ requires only the knowledge of how critical force varies with salt concentrations (Δ*G* = 0). For any given salt concentration, the best statistics is obtained at the critical force because at *f*_c_, multiple transitions between the two states can be observed.

From the coefficients in the dependence of *f*_c_ on log *C*_m_, we can estimate ΔΓ which provides a quantitative estimation of the extent of ion-RNA interactions [45]. For the E⇌I transition, (Δ*R*_ee_) ≈ 4.9 nm and the slope of the linear function is 1.7 pN, which leads to ΔΓ_EI_ = 0.96 (Fig. 8C). This indicates that upon the conformational change from unfolded to the intermediate state, one ion pair enters the atmosphere of the PK. For the transition between I and F, fitting the dependence of 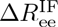 on 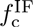 with a = 0.14 nm/pN and b = 0.27 nm (Fig. 9C), we estimate ΔΓ_IF_ = 1.8 using Eq. 6 (Fig. 8D). Consequently, the total number of excess cations effectively bound to folded RNA is expected to be around 2.8, a value that is in rough accord with the experimental estimate of 2.2 [30]. More importantly, ΔΓ_*α*β_ can be measured using LOT and magnetic tweezer experiments [39–43], thus expanding the scope of such experiments to measure ion-RNA interactions. A recent illustration of the efficacy of single-molecule experiments is the study by Jacobson and Saleh who quantified ΔΓ from force measurements for an RNA hairpin [43].

Our analysis is based on the assumption that ΔΓ is a constant independent of the salt concentration. However, it is known experimentally that ΔΓ deviates from a constant value at high salt concentration [41, 43, 46].

In Fig. 8C and D, theoretical lines (blue dotted) show such a deviation from Δ*G* = 0 of the simulation data (white region) at high salt concentration *C* ≳ 500mM). Therefore, we believe that our theoretical analysis is most accurate for *C* ≲ 500 mM. Our coarse-grained simulation reflects the nonlinearity of ΔΓ observed in previous experiments. Understanding the origin of the nonlinearity requires a microscopic theory that should take into account the effects of ion-ion correlations. For complex architectures such as PK or ribozymes, this does not appear straightforward.

## MODEL AND METHODS

### The three-interaction-site (TIS) model

We employed a variant of the three-interaction-site (TIS) model, a coarse-grained model first introduced by Hyeon and Thirumalai for simulating nucleic acids [34]. The TIS model has been previously used to make several quantitative predictions for RNA molecules, ranging from hairpins to ribozymes with particular focus on folding and response to *f* [34, 38, 47–49]. More recently, we have incorporated the consequences of counter ion condensation into the TIS model, allowing us to predict thermodynamic properties of RNA hairpins and PK that are in remarkable agreement with experiments [37]. In the TIS model, each nucleotide is represented by three coarse-grained spherical beads corresponding to phosphate (P), ribose sugar (S), and a base (B). Briefly, the effective potential energy (for details, see Ref. [37]) of a given RNA conformation is *U*_TIS_ = *U*_L_ + *U*_EV_ + *U*_ST_ + *U*_HB_ + *U*_EL_, where *U*_L_ accounts for chain connectivity and angular rotation of the polynucleic acids, *U*_EV_ accounts for excluded volume interactions of each chemical group, and *U*_ST_ and *U*_HB_ are the base-stacking and HB interactions, respectively. Finally, the term *U*_EL_ corresponds to electrostatic interactions between phosphate groups.

The determination of the parameters for use in simulations of the coarse-grained model of RNA was described in detail in a previous publication [37]. Briefly, we determined the stacking interactions by a learning procedure, which amounts to reproducing the measured melting temperature of dimers. We showed that a single choice of model parameters (stacking interactions, the Debye-HÜckel potential for electrostatic interactions, and structure-specific choice for hydrogen bonds) is sufficient to obtain detailed agreements with available experimental data for three different RNA molecules. Thus, the parameters are transferable and we use them verbatim in the present simulations. The determination of hydrogen bond interaction is predicated on the structure. Most of these are taken from the A-form RNA structure and hence can be readily used for any RNA molecule. Only the interaction lengths and angles in hydrogen-bonding interaction of noncanonical base pairs are based on the specific PDB structure, which in our case is the BWYV pseudoknot in this study. We list all the BYWV-PK-specific values used in our simulations in Table S1.

The repulsive electrostatic interactions between the phosphate groups is taken into account through the Debye-HÜckel theory 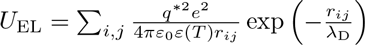, where the Debye length is 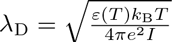. In the simulations, salt concentration can be varied by changing the ionic strength 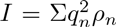 where q_n_ is the charge of ion of type *n*, and ρ_n_ is its number density. Following our earlier study [37], we use an experimentally fitted function for the temperature-dependent dielectric constant ɛ(*T*) [50]. The renormalized charge on the phosphate group is −q\**e*(q* < 1). Because of the highly charged nature of the polyanion, counterions condense onto the RNA, thus effectively reducing the effective charge per phosphate. The extent of charge reduction can be calculated using the Oosawa-Manning theory. Charge renormalization, first used in the context of RNA folding by Heilman-Miller et al. [51] and more recently incorporated into CG models for RNA by us [37] and others [52, 53], is needed to predict accurately the thermodynamics and kinetics of RNA folding [54]. The renormalized value of the charge on the P group is approximately given by 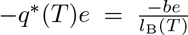, where the Bjerrum length is 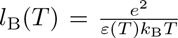 and b is the mean distance between the charges on the phosphate groups [55]. The mean distance (b) between charges on RNA is difficult to estimate (except for rod-like polyelectrolytes) because of its shape fluctuations. Therefore, it should be treated as an adjustable parameter. In our previous study, we showed that b = 4.4 Å provides an excellent fit of the thermodynamics of RNA hairpins and the MMTV pseudoknot [37]. We use the same value in the present study of BWYV as well. Thus, the force field used in this study is the same as in our previous study attesting to its transferability for simulating small RNA molecules.

The native structure of the BWYV PK is taken from the PDB (437D) [32]. The structure contains 28 nucleotides. We added an additional guanosine monophosphate to both the 5’ and 3’ terminus. Thus, the simulated PK has 30 nucleotides. The mechanical force is applied to the ribose sugars of both the 5’– and 3’– terminus (Fig. 1A).

## Simulation details

We performed Langevin dynamics simulations by solving the equation of motion, 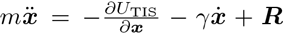, where m is mass; γ, the friction coefficient, depends on the size of the particle type (P, S, or B); and ***R*** is a Gaussian random force, which satisfies the fluctuation-dissipation relation. The [*C*, *f*] phase diagram is calculated at a somewhat elevated temperature, *T*γ50°C, in order to observe multiple folding-unfolding transitions when the monovalent salt concentration is varied over a wide range, from 5 to 1200 mM. The numerical integration is performed using the leap-frog algorithm with time step length δ*t*_L_ δ 0.05τ, where τ is the unit of time. We performed low friction Langevin dynamics simulations to enhance the rate of conformational sampling [56]. For each condition (*C* and *f*), we performed 50 independent simulations for 6 * 10^8^ time steps and the last two-third of trajectories are used for data analyses. In addition, we also performed a set of simulations using the replica-exchange molecular dynamics (REMD) to confirm the validity of the results obtained by the low friction Langevin dynamics simulations [57]. In the REMD simulations, both *C* and *f* are treated as replica variables and are exchanged between 256 replicas, by covering *C* from 1 to 1200 mM and *f* from 0 to 20 pN. The replicas are distributed over 16 * 16 grid. The combined set of simulations using different protocols assures us that the simulations are converged at all conditions.

Our simulations are done at constant forces under equilibrium conditions in which we observe multiple transitions between the various states. We have made comparison to LOT experiments, which are performed by pulling the RNA at a constant velocity. In general, if RNA is stretched at a constant velocity, then one has to be concerned about non-equilibrium effects. However, in optical tweezer experiments, the pulling speed is low enough so that the tension propagates uniformly throughout the RNA prior to initiation of unfolding [58]. Thus, the unfolding of PK studied using LOT experiments [19] occurs at equilibrium, justifying comparisons to our explicit equilibrium simulations.

## Analysis of simulation data

To analyze the structural transitions from which the phase diagrams are determined, we stored snapshots every 10^4^ time steps from all the trajectories. We computed the radius of gyration (*R*_g_) and the end-to-end distance (*R*_ee_) using the stored snapshots. The fraction of native contacts, *Q*, is calculated by counting the number of hydrogen bond (HB) interaction pairs. The assessment of HB formation is based on the instantaneous value of the HB interaction energy, *U*_HB_. In our model, each HB interaction pair contributes up to *n*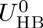 toward stability, where 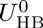 is −2.43 kcal/mol, corresponding to the stability of one HB, and *n* represents number of HB associated in the interaction [37]. A cutoff value *n*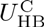 is defined to pick out contacts that are established. We use a value, 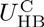 = −1.15 kcal/mol to obtain the diagram using *Q* as an order parameter. Modest changes in 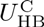 do not affect the results qualitatively.

## CONCLUDING REMARKS

Motivated by the relevance of force-induced transitions in PK to PRF, we have conducted extensive simulations of BWYV PK using a model of RNA, which predicts with near quantitative accuracy the thermodynamics at *f* = 0. The phase diagram in the [*C*, *f*] plane, which can be measured using single-molecule LOT pulling experiments, shows that the thermodynamics of rupture as *f* is increased, or the assembly of the PK as *f* is decreased, involves transitions between the extended, intermediate, and the folded states. The predicted linear relationship between *f*_c_ and log *C*_m_ [Eq. 4] or 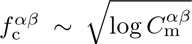 [eq. 6] shows that ΔΓ can be directly measured in LOT experiments. The theoretical arguments leading to Eqs. 4 and (6) are general. Thus, *f* can be used as a perturbant to probe ion-RNA interactions through direct measurement of ΔΓ.

## ACKNOWLEDGMENTS

This work was completed when the authors were at the University of Maryland. We are grateful to Jon Dinman, Xin Li, Pavel Zhuravlev, Huong Vu, and Mauro Mugnai for comments on the manuscript. This work was supported in part by a grant from the National Science Foundation (CHE–1636424).

---

**FIG. S1.**
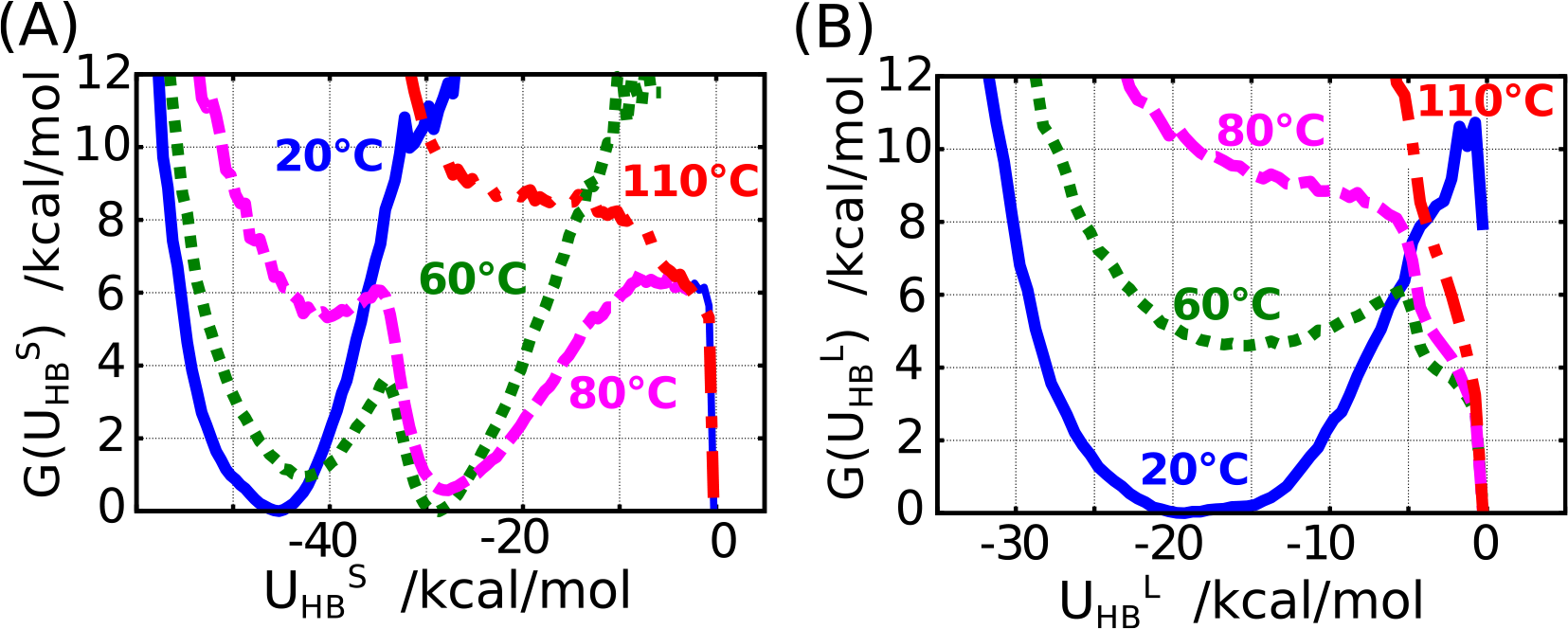
Free energy profiles of the hydrogen bond energy of two stems (A; 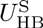) and two loops (stem-loop interactions) (B; 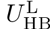) at *C* = 500 mM and *f* = 0.

**FIG. S2.**
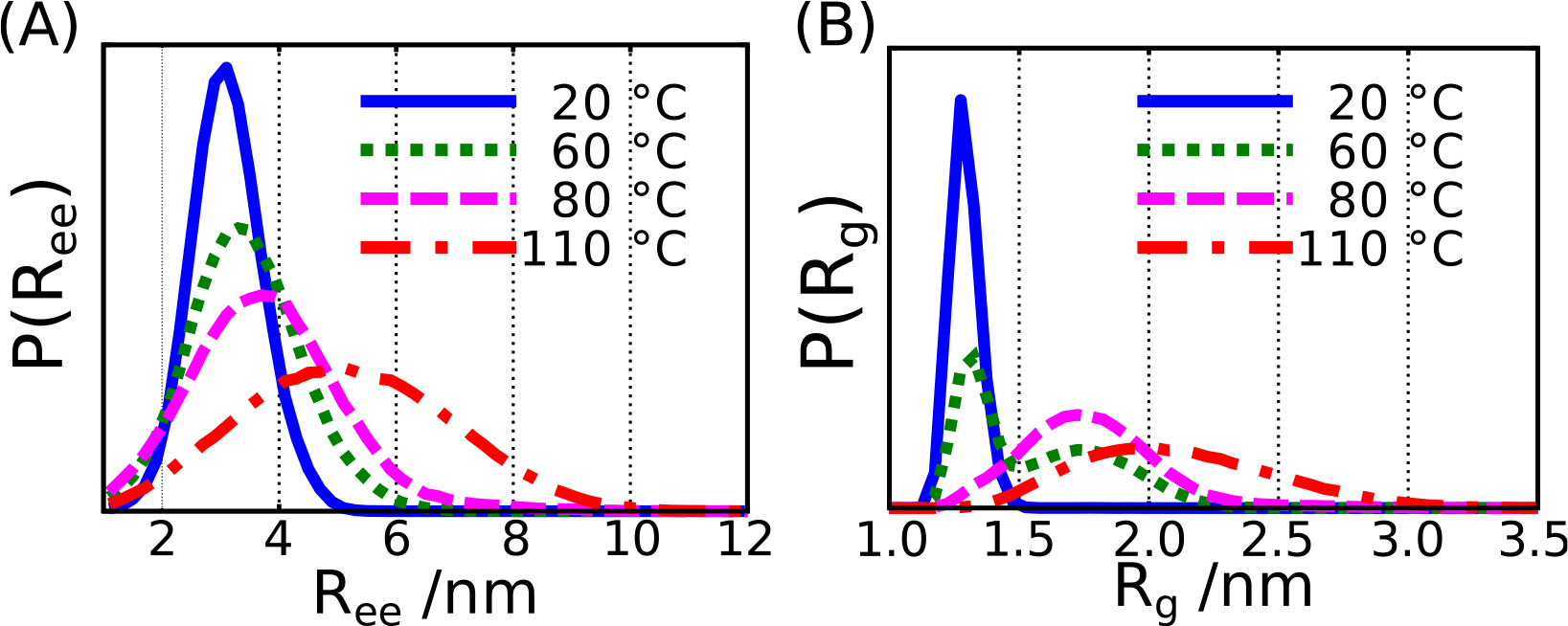
Probability distributions of the molecular extension (A; *R*_ee_) and radius of gyration (B; *R*_g_) at *C* = 500 mM and *f* = 0.

**FIG. S3.**
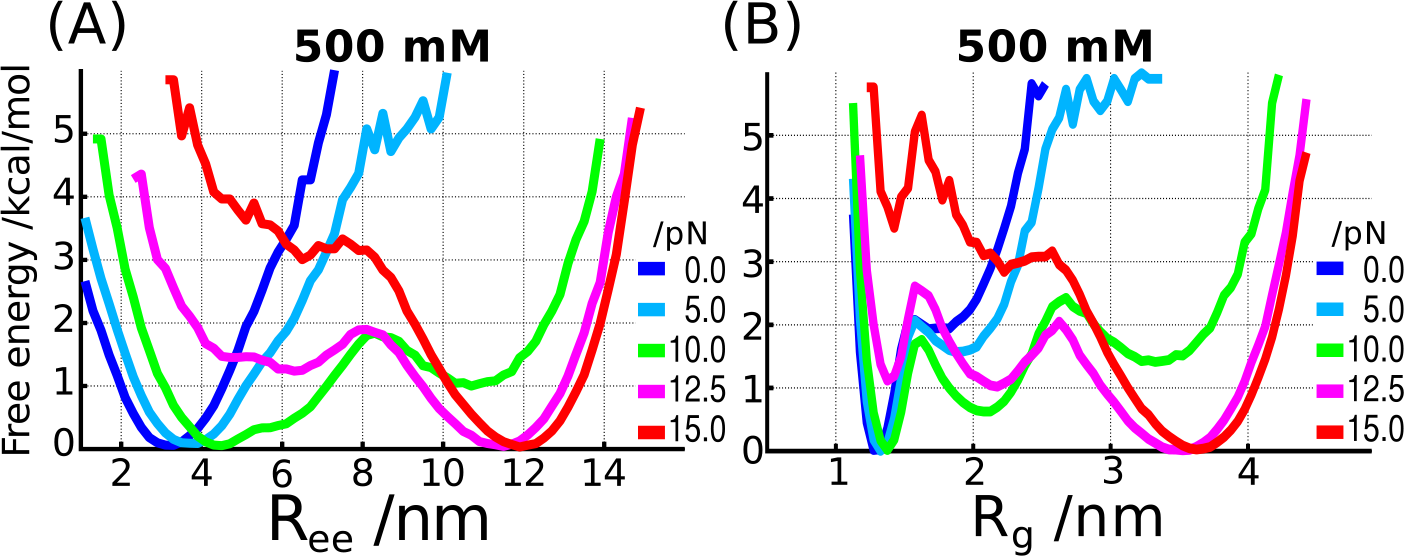
Free energy profiles as a function of *R*_ee_ (A) and *R*_g_ (B) at 500 mM of monovalent salt.

**FIG. S4.**
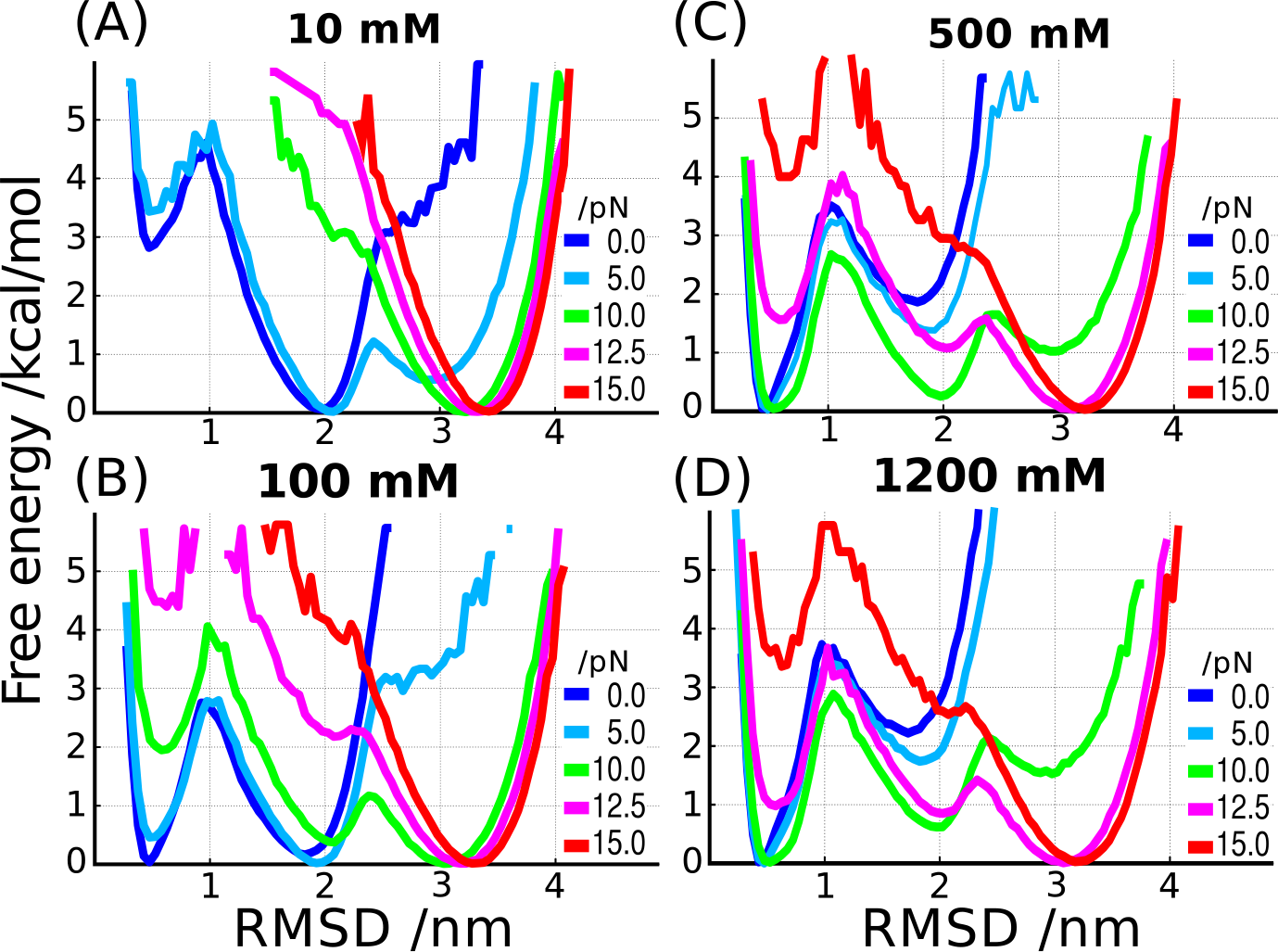
Free energy profiles as a function of RMSD at four different salt concentrations, 10 mM (A), 100 mM (B), 500 mM (C), and 1200 mM (D).

**FIG. S5.**
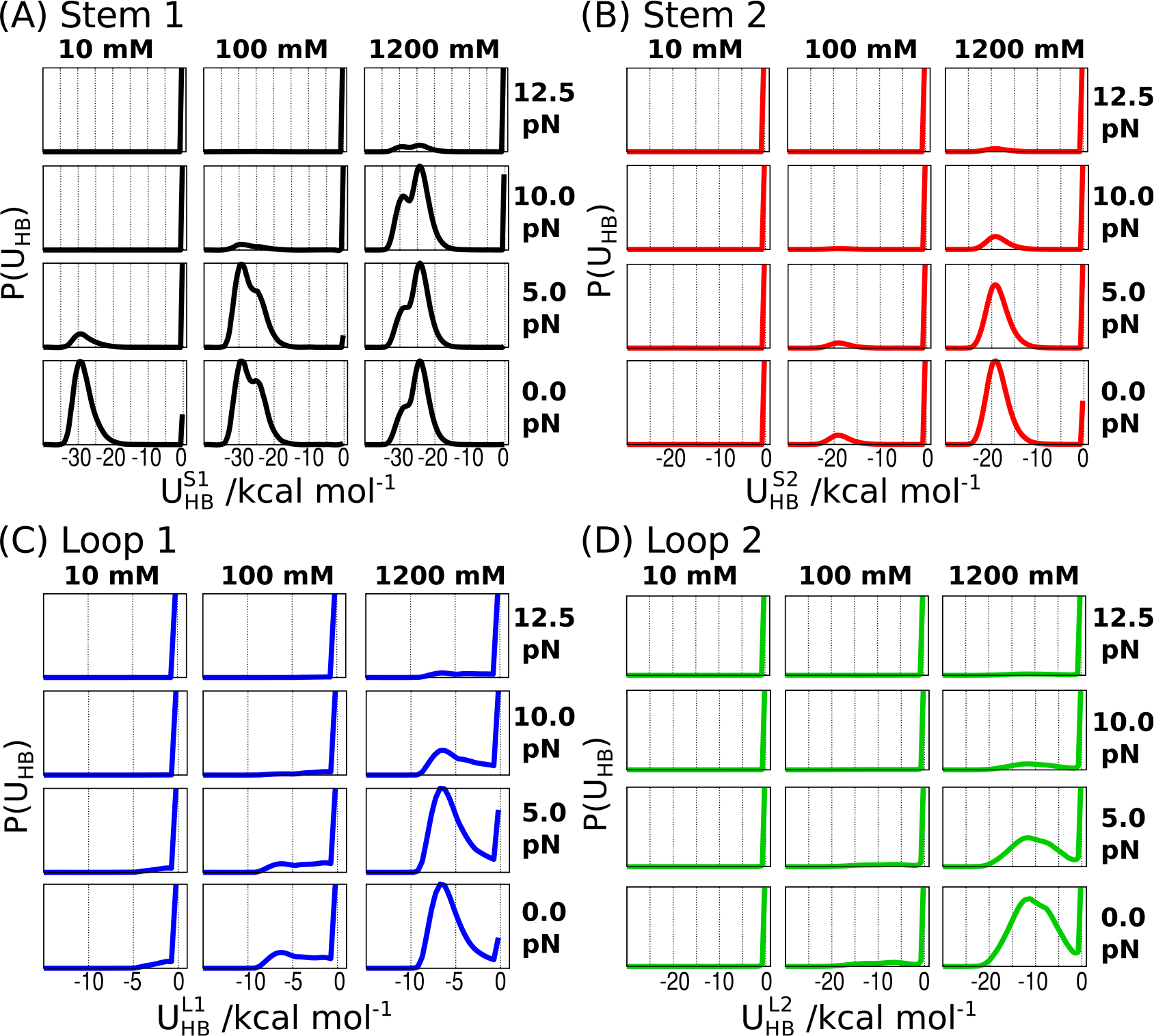
Probability distributions of hydrogen-bond energy. (A-D) Distributions of hydrogen bond energies for individual structural elements (A) Stem1; (B) Stem 2; (C) Loop 1; and (D) Loop2. The salt concentrations and force values are explicitly indicated.

**Table S1.**
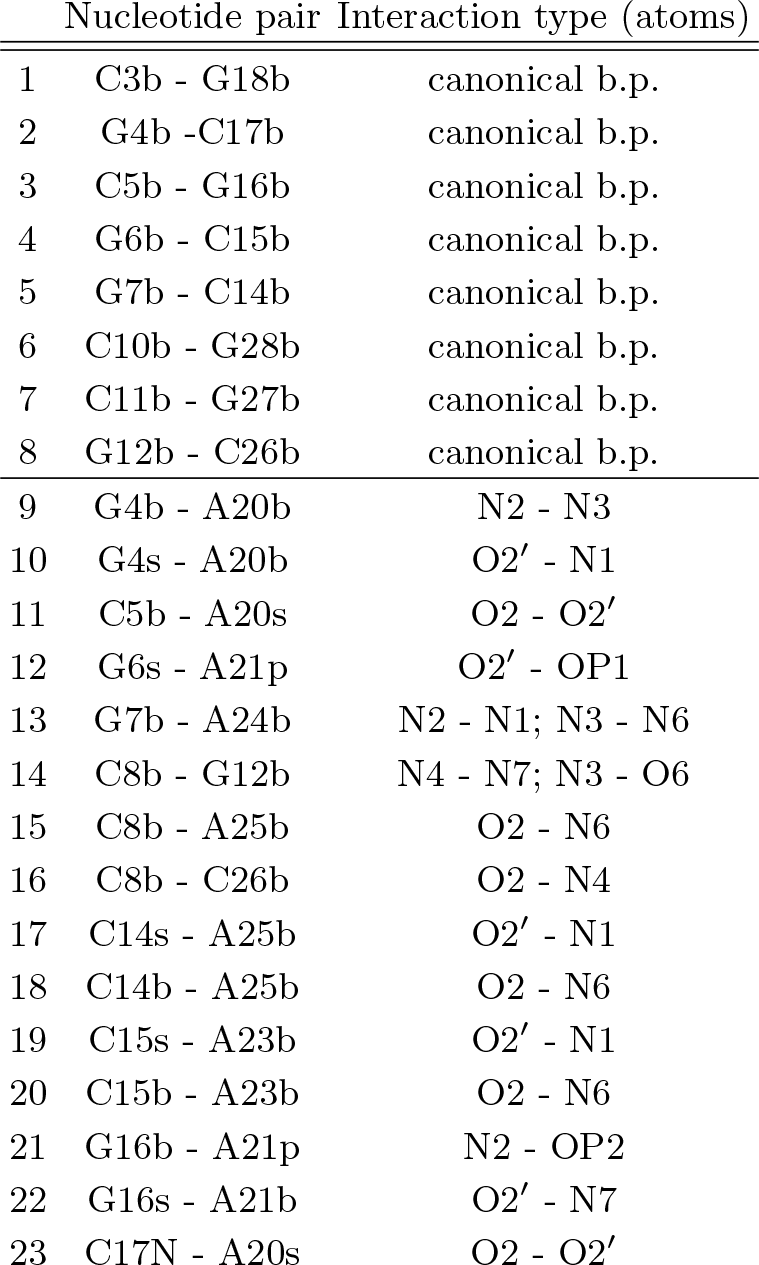
List of hydrogen bonding interactions in BWYV PK. First 8 pairs correspond canonical base pairs forming stems S1 and S2. Other 15 pairs are all tertially interactions forming either L1 or L2. Since we use the three-interaction-site model, a letter “s”, “b” or “p” is put following each nucleotide number to indicate one of sugar, base, or phosphate, respectively.

## References

[1] Cech, T. R.; Steitz, J. A. The noncoding RNA revolution—trashing old rules to forge new ones. Cell 2014, 157, 77–94, DOI: 10.1016/j.cell.2014.03.008.

[2] Tinoco, I.; Bustamante, C. How RNA folds. J. Mol. Biol. 1999, 293, 271–281, DOI: 10.1006/jmbi.1999.3001.

[3] Woodson, S. A. Structure and assembly of group I introns. Curr. Opin. Struct. Biol. 2005, 15, 324–330, DOI: 10.1016/j.sbi.2005.05.007.

[4] Thirumalai, D.; Hyeon, C. RNA and protein folding: common themes and variations. Biochemistry 2005, 44, 4957–70, DOI: 10.1021/bi047314.

[5] Chen, S.-J. RNA folding conformational statistics, folding kinetics, and ion electrostatics. Annu. Rev. Biophys. 2008, 37, 197, DOI: 10.1146/annurev.biophys.37.032807.125957.

[6] Woodson, S. A. RNA folding pathways and the self-assembly of ribosomes. Acc. Chem. Res. 2011, 44, 1312–1319, DOI: 10.1021/ar2000474.

[7] Tinoco, I.; TX Li, P.; Bustamante, C. Determination of thermodynamics and kinetics of RNA reactions by force. Q. Rev. Biophys. 2006, 39, 325–360, DOI: 10.1017/S0033583506004446.

[8] Theimer, C. A.; Feigon, J. Structure and function of telomerase RNA. Curr. Opin. Struct. Biol. 2006, 16, 307–18, DOI: 10.1016/j.sbi.2006.05.005.

[9] Brierley, I.; Digard, P.; Inglis, S. C. Characterization of an efficient coronavirus ribosomal frameshifting signal: requirement for an RNA pseudoknot. Cell 1989, 57, 537–547, DOI: 10.1016/0092-8674(89)90124-4.

[10] Powers, T.; Noller, H. F. A functional pseudoknot in 16S ribosomal RNA. EMBO J. 1991, 10, 2203.

[11] ten Dam, E.; Pleij, K.; Draper, D. Structural and functional aspects of RNA pseudoknots. Biochemistry 1992, 31, 11665–11676, DOI: 10.1021/bi00162a001.

[12] Staple, D. W.; Butcher, S. E. Pseudoknots: RNA structures with diverse functions. PLoS Biol. 2005, 3, e213, DOI: 10.1371/journal.pbio.0030213.

[13] Gilley, D.; Blackburn, E. H. The telomerase RNA pseudoknot is critical for the stable assembly of a catalytically active ribonucleoprotein. Proc. Natl. Acad. Sci. USA 1999, 96, 6621–6625, DOI: 10.1073/pnas.96.12.6621.

[14] Green, L.; Kim, C.-H.; Bustamante, C.; Tinoco, I. Characterization of the mechanical unfolding of RNA pseudoknots. J. Mol. Biol. 2008, 375, 511–528, DOI: 10.1016/j.jmb.2007.05.058.

[15] Ritchie, D. B.; Foster, D. A. N.; Woodside, M. T. Programmed —1 frameshifting efficiency correlates with RNA pseudoknot conformational plasticity, not resistance to mechanical unfolding. Proc. Natl. Acad. Sci. USA 2012, 109, 16167–16172, DOI:10.1073/pnas.1204114109.

[16] Kim, H.-K.; Liu, F.; Fei, J.; Bustamante' C.; Gonzalez, R. L.; Tinoco, I. A frameshifting stimulatory stem loop destabilizes the hybrid state and impedes ribosomal translocation. Proc. Natl. Acad. Sci. USA 2014, 111, 5538–5543, DOI: 10.1073/pnas.1403457111.

[17] Chen, G.; Wen, J.-D.; Tinoco, I. Single-molecule mechanical unfolding and folding of a pseudoknot in human telomerase RNA. RNA 2007, 13, 2175–2188, DOI: 10.1261/rna.676707.

[18] Giedroc, D. P.; Cornish, P. V. Frameshifting RNA pseudoknots: structure and mechanism. Virus Res. 2009, 139, 193–208, DOI: 10.1016/j.virusres.2008.06.008.

[19] White, K. H.; Orzechowski, M.; Fourmy, D.; Visscher, K. Mechanical unfolding of the beet western yellow virus -1 frameshift signal. J. Am. Chem. Soc. 2011, 133, 9775–9782, DOI: 10.1021/ja111281f.

[20] Dinman, J. D. Mechanisms and implications of programmed translational frameshifting. WIREs RNA 2012, 3, 661–673, DOI: 10.1002/wrna.1126.

[21] Chen, G.; Chang, K.-Y.; Chou, M.-Y.; Bustamante, C.; Tinoco, I. Triplex structures in an RNA pseudoknot enhance mechanical stability and increase efficiency of -1 ribosomal frameshifting. Proc. Natl. Acad. Sci. USA 2009, 106, 12706–11, DOI: 10.1073/pnas.0905046106.

[22] de Messieres, M.; Chang, J.-C.; Belew, A. T.; Meskauskas, A.; Dinman, J. D.; La Porta, A. Singlemolecule measurements of the CCR5 mRNA unfolding pathways. Biophys. J. 2014, 106, 244–252, DOI: 10.1016/j.bpj.2013.09.036.

[23] Napthine, S.; Liphardt, J.; Bloys, A.; Routledge, S.; Brierley, I. The role of RNA pseudoknot stem 1 length in the promotion of efficient -1 ribosomal frameshifting. J. Mol. Biol. 1999, 288, 305–320, DOI: 10.1006/jmbi.1999.2688.

[24] Kontos, H.; Napthine, S.; Brierley, I. Ribosomal pausing at a frameshifter RNA pseudoknot is sensitive to reading phase but shows little correlation with frameshift efficiency. Mol. Cell Biol. 2001, 21, 8657–8670, DOI: 10.1128/MCB.21.24.8657-8670.2001.

[25] Kim, Y.-G.; Su, L.; Maas, S.; O’Neill, A.; Rich, A. Specific mutations in a viral RNA pseudoknot drastically change ribosomal frameshifting efficiency. Proc. Natl. Acad. Sci. USA 1999, 96, 14234–14239, DOI: 10.1073/pnas.96.25.14234.

[26] Mouzakis, K. D.; Lang, A. L.; Vander Meulen, K. A.; Easterday, P. D.; Butcher, S. E. HIV-1 frameshift efficiency is primarily determined by the stability of base pairs positioned at the mRNA entrance channel of the ribosome. Nucleic Acids Res. 2013, 41, 1901–1913, DOI: 10.1093/nar/gks1254.

[27] Belew, A. T.; Meskauskas, A.; Musalgaonkar, S.; Advani, V. M.; Sulima, S. O.; Kasprzak, W. K.; Shapiro, B. A.; Dinman, J. D. Ribosomal frameshifting in the CCR5 mRNA is regulated by miRNAs and the NMD pathway. Nature 2014, 512, 265–269, DOI: 10.1038/na-ture13429.

[28] Li, Y.; Treffers, E. E.; Napthine, S.; Tas, A.; Zhu, L.; Sun, Z.; Bell, S.; Mark, B. L.; van Veelen, P. A.; van Hemert, M. J.; Firthe, A. E.; Brierleye, I.; Snijderc, E. J.; Fang, Y. Transactivation of programmed ribosomal frameshifting by a viral protein. Proc. Natl. Acad. Sci. USA 2014, 111, E2172–E2181, DOI: 10.1073/pnas.1321930111.

[29] Tionoco, I.; Bustamante, C. The effect of force on thermodynamics and kinetics of single molecule reactions. Biophys. Chem. 2002, 101, 513–533, DOI: 10.1016/S0301-4622(02)00177-1.

[30] Nixon, P. L.; Giedroc, D. P. Energetics of a strongly pH dependent RNA tertiary structure in a frameshifting pseudoknot. J. Mol. Biol. 2000, 296, 659–71, DOI: 10.1006/jmbi.1999.3464.

[31] Soto, A. M.; Misra, V.; Draper, D. E. Tertiary structure of an RNA pseudoknot is stabilized by “diffuse” Mg^2+^ ions. Biochemistry 2007, 46, 2973–83, DOI: 10.1021/bi0616753.

[32] Su, L.; Chen, L.; Egli, M.; Berger, J. M.; Rich, A. Minor groove RNA triplex in the crystal structure of a ribosomal frameshifting viral pseudoknot. Nat. Struct. Mol. Biol. 1999, 6, 285–292, DOI: 10.1038/6722.

[33] Tan, Z.-J.; Chen, S.-J. Salt contribution to RNA tertiary structure folding stability. Biophys. J. 2011, 101, 176–87, DOI: 10.1016/j.bpj.2011.05.050.

[34] Hyeon, C.; Thirumalai, D. Mechanical unfolding of RNA hairpins. Proc. Natl. Acad. Sci. USA 2005, 102, 6789–94, DOI: 10.1073/pnas.0408314102.

[35] Anderson, C. F.; Record Jr, M. T. Salt dependence of oligoion-polyion binding: a thermodynamic description based on preferential interaction coefficients. J. Phys. Chem. 1993, 97, 7116–7126, DOI: 10.1021/j100129a032.

[36] Liu, T.; Kaplan, A.; Alexander, L.; Yan, S.; Wen, J.-D.; Lancaster, L.; Wickersham, C. E.; Fredrick, K.; Noller, H.; Tinoco, I.; Bustamante, C. J. Direct measurement of the mechanical work during translocation by the ribosome. eLife 2014, 3, e03406, DOI: 10.7554/eLife.03406.

[37] Denesyuk, N. A.; Thirumalai, D. Coarse-grained model for predicting RNA folding thermodynamics. J. Phys. Chem. B 2013, 117, 4901–11, DOI: 10.1021/jp401087x.

[38] Cho, S. S.; Pincus, D. L.; Thirumalai, D. Assembly mechanisms of RNA pseudoknots are determined by the stabilities of constituent secondary structures. Proc. Natl. Acad. Sci. USA 2009, 106, 17349–54, DOI: 10.1073/pnas.0906625106.

[39] Todd, B. A.; Rau, D. C. Interplay of ion binding and attraction in DNA condensed by multivalent cations. Nucleic Acids Res. 2007, 36, 501–510, DOI: 10.1093/nar/gkm1038.

[40] Zhang, H.; Marko, J. F. Maxwell relations for single-DNA experiments: Monitoring protein binding and double-helix torque with force-extension measurements. Phys. Rev. E 2008, 77, 031916–9, DOI: 10.1103/Phys-RevE.77.031916.

[41] Dittmore, A.; Landy, J.; Molzon, A. A.; Saleh, O. A. Single-molecule methods for ligand counting: Linking ion uptake to DNA hairpin folding. J. Am. Chem. Soc. 2014, 136, 5974–5980, DOI: 10.1021/ja500094z.

[42] Jacobson, D. R.; Saleh, O. A. Measuring the differential stoichiometry and energetics of ligand binding to macromolecules by single-molecule force spectroscopy: an extended theory. J. Phys. Chem. B 2015, 119, 1930–1938, DOI: 10.1021/jp511555g.

[43] Jacobson, D. R.; Saleh, O. A. Quantifying the ion atmosphere of unfolded, single-stranded nucleic acids using equilibrium dialysis and single-molecule methods. Nucleic Acids Res. 2016, 44, 3763–3771, DOI: 10.1093/nar/gkw196.

[44] Bond, J. P.; Anderson, C. F.; Record Jr, M. T. Conformational transitions of duplex and triplex nucleic acid helices: thermodynamic analysis of effects of salt concentration on stability using preferential interaction coefficients. Biophys. J. 1994, 67, 825, DOI: 10.1016/S0006-3495(94)80542-9.

[45] Record Jr, M. T.; Zhang, W.; Anderson, C. F. Analysis of effects of salts and uncharged solutes on protein and nucleic acid equilibria and processes: a practical guide to recognizing and interpreting polyelectrolyte effects, Hofmeister effects, and osmotic effects of salts. Adv. Protein Chem 1998, 51, 281–353.

[46] Record Jr, M. T.; Anderson, C. F.; Lohman, T. M. Thermodynamic analysis of ion effects on the binding and conformational equilibria of proteins and nucleic acids: the roles of ion association or release, screening, and ion effects on water activity. Q. Rev. Biophys. 1978, 11, 103–178, DOI: 10.1017/S003358350000202X.

[47] Biyun, S.; Cho, S. S.; Thirumalai, D. Folding of human telomerase RNA pseudoknot using ion-jump and temperature-quench simulations. J. Am. Chem. Soc. 2011, 133, 20634–43, DOI: 10.1021/ja2092823.

[48] Lin, J.-C.; Thirumalai, D. Kinetics of allosteric transitions in S-adenosylmethionine riboswitch are accurately predicted from the folding landscape. J. Am. Chem. Soc. 2013, 135, 16641–50, DOI: 10.1021/ja408595e.

[49] Denesyuk, N. A.; Thirumalai, D. How do metal ions direct ribozyme folding? Nature Chem. 2015, 7, 793–801, DOI: 10.1038/nchem.2330.

[50] Malmberg, C. G.; Maryott, A. A. Dielectric constant of water from 0° to 100° C. J. Res. Nat. Bur. Stand. 1956, 56.

[51] Heilman-Miller, S. L.; Thirumalai, D.; Woodson, S. A. Role of counterion condensation in folding of the Tetrahymena ribozyme. I. Equilibrium stabilization by cations. J. Mol. Biol. 2001, 306, 1157–1166, DOI: 10.1006/jmbi.2001.4437.

[52] Hayes, R. L.; Noel, J. K.; Whitford, P. C.; Mohanty, U.; Sanbonmatsu, K. Y.; Onuchic, J. N. Reduced model captures Mg^2+^-RNA interaction free energy of riboswitches. Biophys. J. 2014, 106, 1508–1519, DOI: 10.1016/j.bpj.2014.01.042.

[53] Hayes, R. L.; Noel, J. K.; Mandic, A.; Whitford, P. C.; Sanbonmatsu, K. Y.; Mohanty, U.; Onuchic, J. N. Generalized Manning condensation model captures the RNA ion atmosphere. Phys. Rev. Lett. 2015, 114, 258105, DOI: 10.1103/PhysRevLett.114.258105.

[54] Heilman-Miller, S. L.; Pan, J.; Thirumalai, D.; Woodson, S. A. Role of counterion condensation in folding of the Tetrahymena ribozyme II. Counterion-dependence of folding kinetics. J. Mol. Biol. 2001, 309, 57–68, DOI: 10.1006/jmbi.2001.4660.

[55] Manning, G. Limiting Laws and Counterion Condensation in Polyelectrolyte Solutions I. Colligative Properties. J. Chem. Phys. 1969, 51, 924, DOI: 10.1063/1.1672157.

[56] Honeycutt, J. D.; Thirumalai, D. The nature of folded states of globular proteins. Biopolymers 1992, 32, 695–709, DOI: 10.1002/bip.360320610.

[57] Sugita, Y.; Okamoto, Y. Replica-exchange molecular dynamics method for protein folding. Chem. Phys. Lett. 1999, 314, 141–151, DOI: 10.1016/S0009-2614(99)01123-9.

[58] Hyeon, C.; Dima, R. I.; Thirumalai, D. Pathways and kinetic barriers in mechanical unfolding and refolding of RNA and proteins. Structure 2006, 14, 1633–1645, DOI: 10.1016/j.str.2006.09.002.

